# Widespread convergent evolution of alpha-neurotoxin resistance in African mammals

**DOI:** 10.1101/2022.09.14.508034

**Authors:** Danielle H. Drabeck, Jennifer Holt, Suzanne E. McGaugh

## Abstract

Convergent evolution is central to the study of adaptation and has been used to understand both the limits of evolution and the diverse patterns and processes which result in adaptive change. Resistance to snake venom α-neurotoxins (αNTXs) is a case of widespread convergence having evolved several times in snakes, lizards, and mammals. Despite extreme toxicity of αNTXs, substitutions in its target, the nicotinic acetylcholine receptor (nAChR), prevent αNTX binding and render species resistant. Recently, the published meerkat (Herpestidae) genome revealed that meerkats have the same substitutions in nAChR as the venom resistant Egyptian mongoose (Herpestidae), suggesting that venom-resistant nAChRs may be ancestral to Herpestids. Like the mongoose, many other species of feliform carnivores prey on venomous snakes, though their venom resistance has never been explored. To evaluate the prevalence and ancestry of αNTX resistance in mammals, we generate a dataset of mammalian nAChR utilizing museum specimens and public datasets. We find five instances of convergent evolution within feliform carnivores, and an additional eight instances across all mammals sampled. Tests of selection show that these substitutions are evolving under positive selection. Repeated convergence suggests that this adaptation played an important role in the evolution of mammalian physiology and potentially venom evolution.

## Introduction

Convergent evolution has offered insight into the ways that diverse organisms evolved to cope with similar selective pressures [1-2]. Within mammals, venom resistance has convergently evolved in marsupials, rodents, carnivores, eulypotyphlans, and artiodactyls [3-9]. Old world mammals which prey upon and sustain bites from venomous snakes in the family Elapidae face envenomation with deadly α-neurotoxins (αNTXs) which bind to the muscular nicotinic acetylcholine receptor (nAChR), blocking nerve-muscle communication and causing rapid muscular paralysis and death. Despite this, the Egyptian mongoose, honey badger, domestic pig, and hedgehogs regularly prey on Elapids and have convergently evolved mutations in their nAChRs which confer resistance to α-NTXs and experience strong positive selection [3].

Mutations associated with αNTX resistance in mammals occur at four sites in the nAChR epitope, and function via three distinct biophysical mechanisms [9]. The first two sites 187 and 189, are aromatic amino acids (W^187^, F^189^) in most mammals, and directly interact with αNTX. Resistance at these sites is present either via steric hindrance mediated by a glycosylation (N^187^, T^189^) in mongooses or via arginine mediated electrostatic repulsion (R^187^) in honey badgers, hedgehogs, and pigs. The second two sites involved in mammalian resistance, 194 and 197 are prolines in most mammals, are necessary for αNTX binding, and replacement of either (e.g., P^194^ to L^194^, P^197^ to H^197^ in the Egyptian mongoose) results in loss of αNTX binding [9]. Extensive experimental work has validated the function of these substitutions in diverse genetic backgrounds [10-20]. Hereafter, we refer to these mechanisms as electrostatic repulsion resistance, glycosylation resistance, and proline resistance, respectively.

Other species of mongooses and meerkats in the same family (Herpestidae) are well-known to exhibit lack-of-avoidance, cooperative mobbing, and predation upon venomous snakes (Figure 1, Supplementary Table 3) [7,21]. The newly published meerkat genome reveals that meerkats share the same changes as the Egyptian mongoose (i.e., N^187^, T^189^ and L^194^) and likely enjoy the same protective glycosylation and partial proline-mediated resistance. αNTX resistance in related species of African feliform carnivores remains unexplored, including in the families Eupleridae (Malagassy carnivores) and Viverridae (civet cats), the latter of which contains many species that exhibit predatory and/or aggressive behavior towards venomous snakes (Figure 1, Supplementary Table 3) [22]. While their ecology, biogeography, and relatedness to resistant taxa suggest that snake venom may be an important selective pressure for many species in Herpestidae, Eupleridae, and Viverridae, it is unknown if they are resistant to αNTX, and whether resistance is the result of an ancient adaptation at the base of these three clades or has evolved convergently in response to repeated selection pressure for venom resistance.

**Figure 1.**
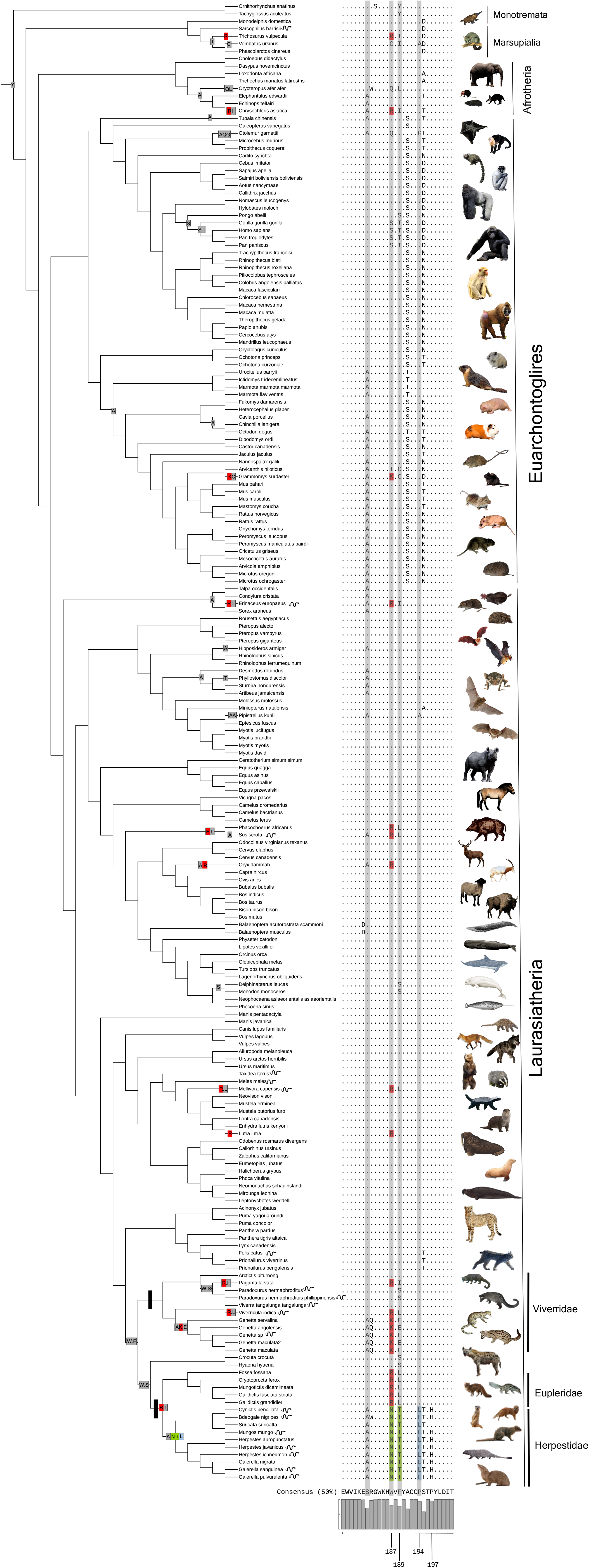
The evolutionary tree of mammals showing the relationships between species for which the nAChR A1 subunit epitope was available. The nAChR epitope alignment from sites 175-198 is displayed for each species with dots denoting amino acids which do not differ from the consensus. Designated foreground clades (Herpestidae, Viverridae, and Eupleridae) are marked with black rectangles. Sites highlighted in gray rectangles indicate signatures of positive selection in these clades (Table 2). Mutations known to confer resistance are reconstructed onto their corresponding branches in color based on the mechanism of resistance: Electrostatic repulsion via a positively charged replacement (R or K) at site 187 is marked in red, steric hindrance via a glycosylation indicated by an N-T replacement at sites 187/189 are marked in green, and replacement of prolines 194 and/or 197 (proline resistance) is marked in blue. Changes at functional and selected sites are mapped onto branches of the tree using a maximum-likelihood based codon model ancestral state reconstruction [29]. Species known to prey on venomous snakes are denoted with a snake icon (citations and summary of interactions in supplementary table 3).

To examine the evolution of venom resistance across these clades, we leveraged an exhaustive search of museum tissues from all species available from North American museums for Eupleridae, Viverridae, and Herpestidae and sequenced the muscular nAChR. We also used bioinformatic searches to identify muscular nAChR sequences from additional species. Using a comparative phylogenetic approach, we leveraged maximum likelihood tests of selection, as well as ancestral sequence reconstruction, to examine whether the species exhibited resistant nAChR, what mechanisms were present, and whether resistance mutations arose ancestrally or convergently across clades.

## Methods

### Tissues and DNA extraction

Thirty-nine tissue samples across 33 different species from Eupleridae, Herpestidae and Viverridae were obtained through specimen loans (Supplementary Table 1). Genomic DNA was extracted as previously described [3] with a skin wash step for recalcitrant samples [23]. Previously designed primers were used to amplify 850 bp of the alpha subunit of the muscular nicotinic acetylcholine receptor gene (*chrna1*) that included the ligand binding (and αNTX) epitope corresponding to residues 122–205 of the protein sequence (See Supplementary Methods). Amplified PCR products were treated with ExoSAP-IT and Sanger sequenced by the University of Minnesota Genomics Center on an ABI 3730XL DNA Analyzer using BigDye Terminator v3.1 chemistry (Applied Biosystems, USA). Resulting DNA sequences were assembled, edited, and aligned using Geneious v8.1 [3, 24].

### Bioinformatic retrieval of sequence

All mammalian sequences included in the NCBI ortholog database (Accessed May 24, 2022) for the gene containing the muscular nAChR sequence, *chrna1*, were included in this dataset. Additional blastP searches were conducted using Viverrid, Herpestid, and Euplerid sequences generated from this project. Results were filtered for duplicate species and a final dataset of 199 sequences was aligned using MUSCLE [24] and edited in Geneious v8.1. A 90bp fragment (175-198aa) covering the acetylcholine/αNTX binding epitope was used for analyses, as this was the longest fragment with complete data for most species (Supplementary Table 3, Supplementary Data). Timetree was used to generate a phylogenetic tree, and missing species were subsequently added in Mesquite using existing topologies [25-28].

### Tests of Selection

We used a suite of likelihood-based codon models which use the ratio of non-silent substitutions to silent substitutions (dN/dS) to detect positive selection [29-35]. Because we were interested in selection specifically on branches of the tree leading to Viverridae, and Herpestidae + Eupleridae (Figure 1), we used a branch-site test for positive selection identifying these lineages and all subtending branches as the foreground. Similarly, site tests were used to identify sites which exhibit signatures of selection [29, 35]. Lastly, to discern signals of convergent versus pervasive selection, we employed a ‘drop out’ site test in which we removed all species in Viverridae, Herpestidae, and Eupleridae as well as honey badger (*Mellivora capensis)*, pig (*Sus scrofa*), and hedgehog (*Erinaceous europea*) from the topology in Mesquite and re-run site tests [26, 36]. Ancestrally reconstructed sequences were generated from the best-fit site model (M8) in PAML v4.6 (Table 1) and aligned using MUSCLE v3.5 [25,29]. For complete details see supplementary methods.

**Table 1.** Results of branch-site tests for positive selection on the CHRNA1 gene.

| Model<br>and log-likelihood | site class<sup>1</sup> | proportion of sites | w background | w foreground<sup>2</sup> |
| --- | --- | --- | --- | --- |
| $w_2 = 1$<br>$\ln L = -1395.62$ | 0 | 0.746 | 0.0139 | 0.0139 |
|  | 1 | 0.152 | 1.0 | 1.0 |
|  | 2a | 0.085 | 0.0139 | 1.0 |
|  | 2b | 0.017 | 1.0 | 1.0 |
| $w_2 > 1$<br>$\ln L = -1384.06$ | site class | proportion of sites | w background | w foreground<sup>2</sup> |
|  | 0 | 0.705 | 0.013 | 0.013 |
|  | 1 | 0.150 | 1.0 | 1.0 |
|  | 2a | 0.121 | 0.013 | 10.20 |
|  | 2b | 0.026 | 1.0 | 10.20 |

<sup>1</sup> Site class 0 and 1 apply to foreground and background lineages and include sites under purifying selection ( $0 < w <$ 1) and neutral sites ( $w = 1$ ), respectively. Site class 2 allows a proportion of positively selected sites ( $w > 1$ ) in the foreground lineages, where 2a includes sites under purifying selection ( $0 < w < 1$ ) in the background lineages and 2b includes neutral sites ( $w = 1$ ) in the background lineages. $2\Delta\ln = 20$ , d.f. =1, $p < 0.0001$ . When *Mellivora capensis*, *Sus* *scrofa*, and *Erinaceus euopus* are included in foreground branches $2\Delta\ln = 30$ , d.f. =1, $p < .00001$
<sup>2</sup> Herpestidae, Eupleridae, and Viverridae were included in the foreground class.

## Results and Discussion

### Evolutionary History of Resistance in Herpestidae, Eupleridae, and Viverridae

To assess αNTX resistance, we used aligned sequence data for the αNTX binding epitope of nAChR (Figure 1) and interpreted sequence data using extensive prior literature that has experimentally demonstrated the effects of specific amino acid substitutions on αNTX binding. We found that all newly sequenced nAChRs from species belonging to the family Herpestidae have N^187^, T^189^ and L^194^, H^197^ which confer strong glycosylation and proline resistance, respectively (Figure 1) [10-20]. All Malagasy carnivores (Eupleridae) as well as two civets (Viverridae) have substitution R^187^ and I/L^189^ which confer electrostatic repulsion resistance (Figure 1) [3,13,18-19]. Within civets (Viverridae), we found novel mutations (K^187^ and E^189^) which arose at the base of the genus *Genetta* (small African carnivores including the genet). Lysine (K) is one of only three positively charged amino acids and may mimic electrostatic repulsion resistance seen with R^187^, however, biophysical testing is needed to definitively assess the impact of K^187^ on αNTX binding.

To examine whether αNTX evolved convergently, we employed a codon model (CODEML) maximum likelihood ancestral reconstruction of sequences implemented in PAML v4.6 [6, 29, 37]. Our ancestral reconstruction supports that electrostatic repulsion resistance via R^187^ arose once at the base of Eupleridae + Herpestidae and twice within Viverridae. Subsequent glycosylation resistance (N^187^, T^189^), as well as Proline resistance (L^194^, H^197^) appears to have arisen later in Herpestidae (Figure 1). Empirical work has shown that electrostatic repulsion resistance is less effective than either glycosylation or proline resistance [9, 12, 20]. Our data suggest that electrostatic repulsion resistance arose first at the base of Eupleridae + Herpestidae, and subsequent selection for increased resistance in Herpestids likely led to glycosylation and proline resistance. Interestingly, the Malagasy carnivores (Euplerids) are not sympatric with any αNTX producing snake besides sea snakes (for which we do not have any evidence of predation). As the Euplerids are restricted to Madagascar, and the R^187^ appears to have preceded the split between Euplerids and Herpestids (Figure 1), the presence of this mutation in these species is most likely a remnant of selection pressure imposed prior to the isolation of this group on Madagascar.

Across the clades Herpestidae, Eupleridae, and Viverridae, mutations known to confer resistance have arisen at least four times (five, if we include K^187^ in civets). These results strongly suggest that species among all three clades have substantial αNTX-resistant nAChRs and warrant further investigation into their ecological interactions with venomous snakes, as well as their physiological ability to cope with venom.

### Positive selection identified for substitutions

To test for adaptive evolution in nAChR in the clades we suspected of being venom resistant, we used codon-model based (CODEML) branch-site tests of positive selection in PAML v4.6 and explored additional methods and results in supplementary materials [29]. Convergent foreground branches were specified as Eupleridae + Herpestidae, the base of Viverridae, and singular branches leading to honey badger, hedgehog, and pig [3]. Because the latter three species and the mongoose have previously been shown to be under positive selection and may inflate the overall signal of selection, we performed branch-site tests with and without these species (Figure 1). In both cases, we recovered a strong signal of selection in foreground lineages (2ΔL= 30, df = 1, p <0.0001; without honey badger, hedgehog, and pig 2ΔL= 20, df = 1, p <0.0001; Table 1). ‘Drop-out’ site tests showed no selection for a tree pruned of all lineages suspected of being under selection, (2ΔL= 2.4, df = 1, p =0.121), indicating that the signal of positive selection can be attributed to the foreground lineages [36].

Bayes Empirical Bayes (BEB) tests identified four sites to be evolving under positive selection in foreground lineages (Table 2). Of these, three (187, 189, and 197) modulate αNTX binding in empirical studies [12-16, 19-20]. Site 182 was identified as a site under positive selection, though no functional studies exist for this site (Figure 1 denoted in gray).

**Table 2.** Sites identified to be evolving under positive selection via different maximum likelihood analyses.

| Site | Foreground Specified<sup>1</sup><br>PAML | Foreground Specified<sup>1</sup><br>MEME | Foreground unspecified<br>PAML | Foreground unspecified<br>MEME | Mutations | Function |
| --- | --- | --- | --- | --- | --- | --- |
| 182 | **0.96** | - | - | - | W,Q | Unknown |
| 187* | **1.00** | - | **0.965** | - | R<sup>3</sup>, N<sup>2</sup>, K | Sh<sup>2</sup>/Er<sup>3</sup> |
| 189* | **1.00** | **0.9911** | 0.776 | - | L, I, E, T<sup>3</sup> | Sh<sup>3</sup> |
| 195 | - | - | **0.990** | **0.955** | L |  |
| 197* | **0.95** | - | - | - | H | Pr<sup>4</sup> |

<sup>2</sup> Reduced binding via steric hindrance [13,16,18,20]
<sup>3</sup> Reduced binding via electrostatic repulsion [18-19]
<sup>4</sup> Reduced binding via proline binding disruption [11,18]
\*Site directly associated with aNTX binding [18]

### Mutations recovered in other mammals

Within our 199 mammal dataset we found five new instances of independently evolved substitutions known to confer electrostatic repulsion resistance (R^187^) outside the three clades initially examined. In two of these instances, the Eurasian otter (*Lutra lutra*) and the oryx antelope (*Oryx dammah*), R^187^ is not secondarily accompanied by the substitution I^189^ or L^189^ (Figure 1). Empirical work has shown that nAChR loses affinity for αNTX with the R^187^ mutation alone, and that addition of the I^189^, or I^189^ by itself does not confer resistance despite its propensity to be paired with R^187^[19]. However, the presence of the single mutant R^187^ in these data suggests that the accompanying mutation I^189^ is likely not a necessary epistatic mutation and raises the possibility that it may have some function in resistance that is apparent *in vivo* that is not recovered *in vitro* [19]. Both the Eurasian otter and the oryx antelope are sympatric with αNTX producing snakes, though direct ecological links were not found in the literature for either species. The Eurasian otter is known to eat venomous catfish (*Ictalurus nebulosus*) which contain nAChR-targeting toxins [38-41]. Further investigations of species closely related to *Oryx dammah* revealed that all available species in the families Alcelaphinae and Hippotraginae also had an R^187^ mutation (Supplementary Material, Supplementary Figure 1).

We found that the warthog, *Phacochoerus africanus*, shares the R^187^ and L^189^ mutation with its pig sister taxa (*Sus scrofa*), and that this mutation arose in an ancestor of these two species. Additional African species with charge resistant mutations are the Cape golden mole (*Chrysochloris asiatica*, R^187^) and the thicket rat (*Grammomys surdaster*; K^187^) and are likely prey of venomous snakes that produce αNTX [42-44].

The only non-African species with R^187^ was the Australian brushtail possum (*Trichosurus vulpecula*). While this species is sympatric with αNTX producing elapids, no specific predator/prey relationship was found in the literature. Brushtail possums are also eucalyptus specialists, and this mutation may be related to coping with the high level of ceramides in Eucalyptus oil, which modulate acetylcholine receptor levels and function [45-46]. Several other species (Wombats, Bush Babies, Aardvarks) showed convergent mutations (K^187^, Q^187^, C^187^) of unknown function, and further biochemical assessment is needed.

## Conclusions

The evolution of resistance to αNTX in mammals has previously been categorized as a relatively rare adaptation only known in six mammals [3]. This work revealed 27 new species with amino acid changes known to cause resistance (experimentally validated across diverse taxa), and an additional 17 species with substitutions that are suspected to result in aNTX resistance [10-20]. In total, our work shows that convergent αNTX resistance has evolved at least 11 times within mammals. Further investigation of fauna that interact with αNTX producing snakes is needed to determine whether αNTX resistance translates to whole venom resistance, and whether these substitutions are the result of present or past ecological interactions with venomous species. Our analyses, along with other recent work, suggest that snake and other animal venoms are a source of strong selection pressure likely facilitated via complex coevolutionary interactions that may be the rule rather than the exception, particularly for animals which share habitat with many venomous snakes [5,19,47].

## Acknowledgements

This work was generously funded by the NIH TREM fellowship (K12GM119955), the UMN UROP, and the Minnesota Herpetological Society. Thanks to the Minnesota Supercomputer Institute and the University of Minnesota Genomic Center. Thanks to the American Museum of Natural History (AMNH), California Academy of Sciences (CAS), Denver Museum of Nature & Science (DMNS), Field Museum of Natural History (FMNH), University of Kansas (KU), Museum of Southwestern Biology (MSB), Berkeley’s Museum of Vertebrate Zoology (MVZ), and Texas Tech University (TTU) for generously providing samples. A special thanks to Rafale Vianna Furer for their work on this project as an undergraduate at Macalester College. We are grateful for the valuable input by Yaniv Brandvain, Sharon Jansa, Emilie Richards, Kyle Keepers, Cathy Rushworth, and Emma Roback. Finally, the authors would like to thank Georgia Hart, Natalie Coles, and Matt Holding whose conversations provided the fodder, curiosity, and motivation for this work.

