## Supplementary Methods for "Widespread convergent evolution of alpha-neurotoxin resistance in African mammals"

*DNA protocols*

Polymerase chain reactions (PCRs) were carried out in 20 µL reactions using 5 µL of DNA extraction product, 1.5 µl of 10 mM ACH_F1 (5'-TGCAGATGGTGACTTTGCCATTGTCAAG-3') primer solution, 1.5 µl of 10 mM ACH_R1 (5'-AGTCTGTGGGCAGGTAGAACACC-3') primer solution, 12.5 µL of AccuPrime (Thermo Fisher Scientific Inc.), and 4.5 µL of deionized water [1]. Reactions were performed for 30 cycles of denaturation at 94 °C for 30 seconds, followed by annealing at 58 °C for 15 seconds, and extension at 72 °C for 90 seconds [1]. Reactions were preceded by a 2 minute denaturation at 94 °C and included a final extension at 72 °C for 7 minutes [1]. ExoSAP reactions were carried out in 6.5 uL reactions using 5 µL of PCR product, 0.5 µL of Exo I, and 1 µL of rSAP (Thermo Fisher Scientific Inc.). ExoSAP reactions were performed at 37 °C for 15 minutes and then enzymes were deactivated at 80 °C for 15 minutes.

*Tests of Selection*

Codon model-based tests of selection are available through a variety of methods [2-8]. No method is without shortcoming, and tests of selection should be treated as hypotheses for the location of functional relevance to be followed up with and/or supported by existing empirical testing of physical function.

Because of the depth of background ecological, physiological, and biochemical mutagenesis work on toxin and venom resistance targets, they are ideal for assessing the performance of statistical tests of selection for similar systems [9-26] To examine and compare results for all methods we used this nACHr dataset as well as another dataset, the von Willebrand Factor alpha 1 subunit in 39 species of Didelphid marsupials, to evaluate the performance of several approaches [1,21]. Both genes were previously found to be targets of selection for toxin resistance and have been subsequently shown to have functionally lost their ability to bind to venom toxins, contributing to adaptive physiological function.

Using these data we tested four methods to determine which were best at identifying selection at sites and branches which are empirically verified to have modified physiological function. We employed 1) a codon model based branch-site test in PAML v4.6 [2] 2) an unrestricted branch-site test, aBSREL (adaptive Branch Site Random Effects likelihood [5] 3) a gene-wide unconstrained selection test (BUSTED) [7] and 4) a site test of positive selection (MEME) [6]. For the nAChR data each site test (Bayes Emperical Bayes and MEME) were run with and without foreground branches specified (Table 2).

Branch-site tests allow the user to designate a 'foreground' which includes lineages thought to be under positive selection. Subsequently, a likelihood ratio test compares two nested models which differ in their allowance of ω (dN/dS) to be elevated in the foreground ($\omega > 1$). Models from PAML [2] and HyPhy [3-8] differ in that PAML restricts background lineage site classes to 0

$> \omega < 1$ or $\omega = 1$, whereas HyPhy allows for the background to have any number of site-classes and tests each branch individually, correcting for multiple tests. aBSREL has been shown to have reduced type 1 error and detect selection in 80-90% of branches with the exception of cases of weak selection, small proportions of sites under positive selection, or along short branches [5]. Because many protein-protein interactions can be altered with small changes of large effect, this leaves many functionally selected targets susceptible to being missed by these methods.

For both datasets, branch site tests in PAML identified foreground branches to be evolving under positive selection (main text, Jansa and Voss 2011), and sites under selection were identified using a Bayes Empirical Bayes Approach. The same two datasets were run in HyPhy v2.5.33 aBSREL and in both instances zero branches were identified to be evolving under positive selection. Gene-wide tests of selection (BUSTED) also identified no positive selection for the vWF dataset ($p = 0.096$) and for the nACHr dataset ($p = 0.5$). For both datasets PAML seems to be the more sensitive test. While the Hyphy unrestricted model tests likely provide a more realistic model, signal being restricted to only a few sites and a few branches may not provide enough degrees of freedom or statistical power to be detected using multiple individual branch testing and subsequent correction.

A comparison of the MEME site test and PAML site test for the nAChR dataset with and without foreground specified showed that PAML outperformed MEME in both instances, identifying four sites under selection, three of which are experimentally confirmed to affect function. In contrast, MEME identified only one site of known functional consequence (Table 1). Similar results were apparent for the vWF dataset (Supplementary Figure 1). PAML identified seven sites under selection which correspond to binding sites as well as sights shown to be associated with loss of venom protein binding [27]. MEME identified 5 of the same sites identified by PAML, and an additional two sites within the venom protein-vWF binding pocket, neither of which have been identified as sites which affect function. MEME also identified two sites outside the binding pocket, interestingly one directly associated with folding and the other located four amino acids downstream from a site known to reduce venom binding in alanine scanning mutagenesis [25,28].

While PAML seems to be more adept at identifying sites that are directly associated with binding, it is possible that MEME results are identifying sites that are also associated with function but have yet to be fully explored. However, based on these data, we conclude that for proteins under selection in which a small proportion of sites are known to affect significant functional change, the PAML branch-sites test is the preferred method.

When selection is convergent across sites or pervasive across a tree, we expect restricted models (PAML) to overestimate convergent selection [29]. We compensate for this by using a dropout site test (as described in the main text). However restricted tests of selection do not rule out selection in the background branches, and some background species exhibited putatively functional mutations as well as ecology and biogeography which may have subjected species selection for venom resistance.

*Further Examination of Bovids*

Subsequent to discovering that the African bovid *Oryx dammah* had a mutation ($R^{187}$) associated with binding loss, we examined this lineage further. Using the entire *chrna1* CDS reference sequence for *Oryx dammah* (XM_040228197.1), we used blastn to blast this sequence against a sampling of available genomes in several major Bovid clades. Resulting scaffolds were downloaded and mapped to the reference (XM_040228197.1) in Geneious v8.1. Amino acid sequence for the epitope is shown in Supplementary Figure 2.

We found that along with Oryx, all available species in the families Alcelaphinae (e.g., wildebeest) and Hippotraginae (e.g. antelopes) also had an $R^{187}$ mutation (Supplementary figure 1). Bovids in the family Caprinae (e.g. goat-antelopes) and Reduncinae (e.g., reedbucks, and allies) had the ancestral aromatic (venom sensitive) residues, while species in Antilopinae (e.g. gazelles, true antelopes) Cephalophinae (duikers), and Aepycerotinae (Impalas) all have mutations $Q^{187}$ and $I^{189}$. These mutations have not been empirically tested for their effect on αNTX binding but are shared with the phylogenetically distant ant-eating aardvark (*Orycteropus afer*) and bushbaby (*Otolemur garnetti*), the latter of which also has a proline resistance replacement at site 194. These data suggest that several African bovids may have evolved some resistance to αNTX, perhaps as a defense against predation or accidental trampling which might result in envenomation.

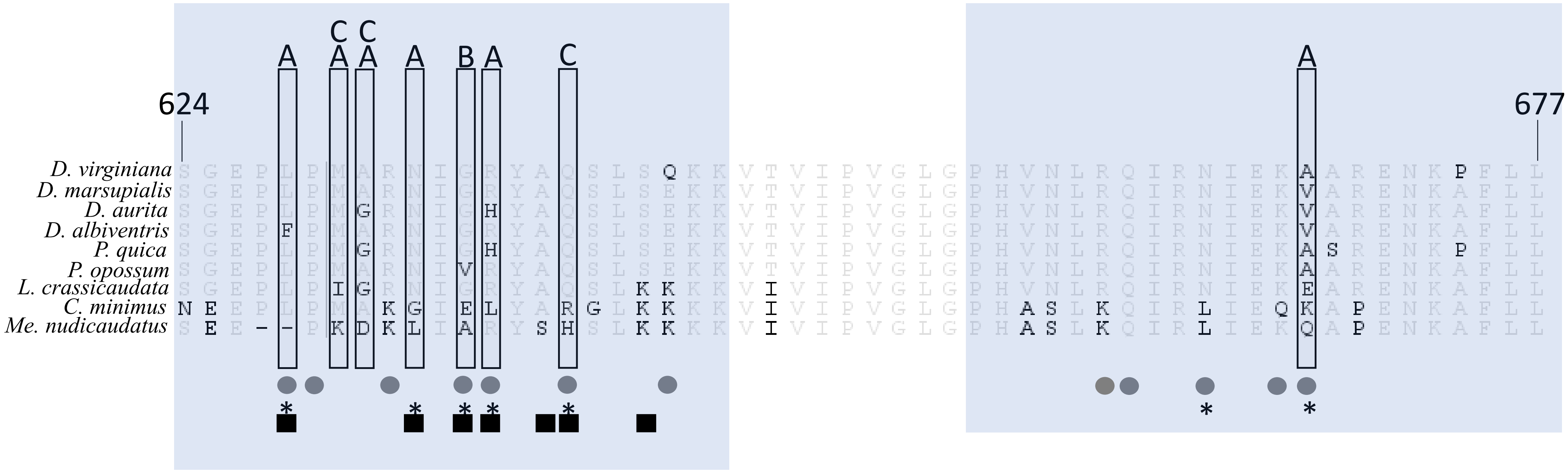

624
677
A
C
B
D. virginiana
D. marsupialis
D. aurita
D. albiventris
P. quica
P. opossum
L. crassicaudata
C. minimus
Me. nudicaudatus

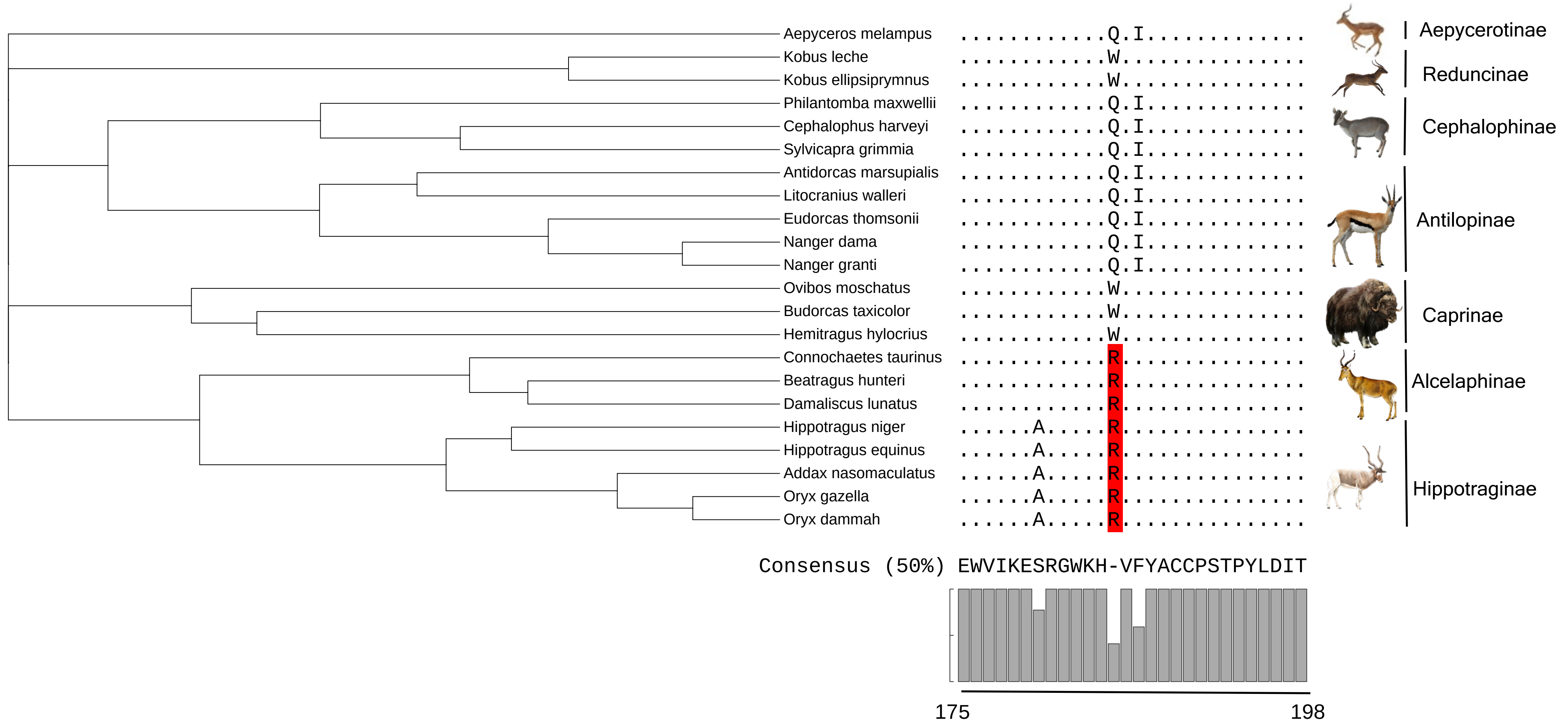

Aepyceros melampus
Kobus leche
Kobus ellipsiprymnus
Philantomba maxwellii
Cephalophus harveyi
Sylvicapra grimmia
Antidorcas marsupialis
Litocranius walleri
Eudorcas thomsonii
Nanger dama
Nanger granti
Ovibos moschatus
Budorcas taxicolor
Hemitragus hylocrius
Connochaetes taurinus
Beatragus hunteri
Damaliscus lunatus
Hippotragus niger
Hippotragus equinus
Addax nasomaculatus
Oryx gazella
Oryx dammah
Aepycerotinae
Reduncinae
Cephalophinae
Antilopinae
Caprinae
Alcelaphinae
Hippotraginae
Consensus (50%) EWVIKESRGWKH-VFYACCPSTPYLDIT
175
198

**Supplementary Figure 1.** Alignment of vWF protein sequences for species of Didelphid opossums (Drabeck et al 2022). Region 624–677 is shown as it encompasses the two amino acid stretches which are known to bind the venom protein botrocetin (624–645) and (655–677) shown in shaded boxes. Majority amino acids are in gray, and variants are denoted in black. Sites associated with *in vitro* binding loss are enclosed in boxes with the venom agonist denoted above the box (A: botrocetin isoform A, B: Botrocetin isoform B, C: aspercetin). Known botrocetin binding sites are marked with a circle. Sites identified as evolving under positive selection by PAML are marked with an asterisk, and sites identified by MEME are marked with a black box.

**Supplementary Figure 2.** The evolutionary tree of bovids from TimeTree showing the relationships between species for which the nAChR A1 subunit epitope was available via NCBI genomes. The nAChR epitope alignment from sites 175-198 is displayed for each species with dots denoting amino acids which do not differ from the consensus. Residues highlighted in red denote mutations which add a positive charge at the binding site (Harris and Fry 2021).

### SUPPLEMENTARY TABLES

**Supplementary Table 1.** Tissue samples queried including Institution, catalog number, family of membership, and locality are listed for all received species' tissue samples. Samples were received from the American Museum of Natural History (AMNH), California Academy of Sciences (CAS), Denver Museum of Nature & Science (DMNS), Field Museum of Natural History (FMNH), University of Kansas (KU), Museum of Southwestern Biology (MSB), Berkeley's Museum of Vertebrate Zoology (MVZ), and Texas Tech University (TTU). Species with superscript were successfully sequenced and accessioned into Genbank (Accession numbers below) and included in the final species alignment.

| Species | Institution | Catalog Number | Family of Membership | Locality | GenBank Accession ID |
|---|---|---|---|---|---|
| *Mungotictis decemlineata* | AMNH | 110067 | Eupleridae | Africa > Madagascar | |
| *Viverra tangalunga tangalunga* | FMNH | 145742 | Viverridae | Asia > Philippine Islands | |
| *Galerella sanguinea* | FMNH | 145231 | Herpestidae | Africa > Uganda | |
| *Mungos mungo* | FMNH | 149364 | Herpestidae | Africa > Burundi | |
| *Galidictis fasciata striata* | FMNH | 156549 | Eupleridae | Africa > | |

| | | | | |
|---|---|---|---|---|
| | | | | Madagascar |
| *Genetta maculata* | FMNH | 149359 | Viverridae | Africa > Burundi |
| *Ichneumia albicauda* | FMNH | 157991 | Herpestidae | Africa > Uganda |
| *Galidictis grandidieri* | FMNH | 173157 | Eupleridae | Africa > Madagascar |
| *Fossa fossana* | FMNH | 156648 | Eupleridae | Africa > Madagascar |
| *Viverricula indica* | FMNH | 187126 | Viverridae | Africa > Tanzania |
| *Genetta tigrina* | FMNH | 177238 | Viverridae | Africa > Mozambique |
| *Paradoxurus hermaphroditus philippinensis* | FMNH | 145743 | Viverridae | Asia > Philippine Islands |
| *Cryptoprocta ferox* | FMNH | 171889 | Eupleridae | Africa > Madagascar |
| *Civettictis civetta* | FMNH | 149356 | Viverridae | Africa > Burundi |
| *Genetta sp. (genetta?)* | FMNH | BDP7276 | Viverridae | ? |
| *Bdeogale nigripes* | FMNH | 167685 | Herpestidae | Africa > Gabon |
| *Genetta angolensis* | FMNH | 163777 | Viverridae | Africa > Tanzania |
| *Bdeogale crassicauda* | FMNH | 208458 | Herpestidae | Africa > Tanzania |
| *Arctictis binturong* | FMNH | 168870 | Viverridae | ? |
| *Helogale parvula* | FMNH | 165387 | Herpestidae | ? |
| *Genetta servalina* | FMNH | 145228 | Viverridae | Africa > Uganda |
| *Proteles cristatus* | DMNS | 5966 | Hyaenidae | Africa > Botswana |
| *Genetta genetta* | DMNS | 5981 | Viverridae | Africa > Botswana |
| *Proteles cristatus* | DMNS | 6008 | Hyaenidae | Africa > Botswana |
| *Hyaena brunnea* | DMNS | 6007 | Hyaenidae | Africa > Botswana |
| *Hyaena brunnea* | DMNS | 9954 | Hyaenidae | Africa > Botswana |

| | | | | |
|---|---|---|---|---|
| *Hyaena brunnea* | DMNS | 6132 | Hyaenidae | Africa > Botswana |
| *Proteles cristatus* | DMNS | 6015 | Hyaenidae | Africa > Botswana |
| *Genetta genetta* | DMNS | 5969 | Viverridae | Africa > Botswana |
| *Genetta genetta* | DMNS | 6021 | Viverridae | Africa > Botswana |
| *Galerella nigrata* | CAS | 32227 | Herpestidae | Africa > Namibia |
| *Galerella pulvurulenta* | CAS | 28740 | Herpestidae | Africa > South Africa |
| *Paradoxurus hermaphroditus* | KU | 171625 | Viverridae | Asia > Philippine Islands |
| *Chrotogale owstoni* | MVZ | 186571 | Viverridae | Asia > Vietnam |
| *Paguma larvata ssp.* | MVZ | 186577 | Viverridae | Asia > Vietnam |
| *Herpestes javanicus* | MSB | 89018 | Herpestidae | United States > Puerto Rico |
| *Crocuta crocuta* | MSB | 281289 | Hyaenidae | United States > New Mexico |
| *Cynictis pencillata* | TTU | 114797 | Herpestidae | Africa > Botswana |
| *Herpestes auropunctatus* | TTU | 43531 | Herpestidae | United States > Puerto Rico |

**Supplementary Table 2**. Results of branch-site tests for positive selection on the CHRNA1 gene with all convergent lineages included.

| Model and log-likelihood | site class[1] | proportion of sites | ω background | ω foreground[2] |
|---|---|---|---|---|
| $\omega_2 = 1$ | 0 | 0.840 | 0.0196 | 0.00196 |
| lnL = -1399.75 | 1 | 0.070 | 1.0 | 1.0 |
| | 2a | 0.083 | 0.0196 | 1.0 |
| | 2b | 0.007 | 1.0 | 1.0 |

| $\omega_2 > 1$ | site class | proportion of sites | ω background | ω foreground[2] |
|---|---|---|---|---|
| lnL = -1384.58 | 0 | 0.694 | 0.012 | 0.012 |
| | 1 | 0.155 | 1.0 | 1.0 |
| | 2a | 0.123 | 0.012 | 7.576 |
| | 2b | 0.027 | 1.0 | 7.576 |

[1] Site class 0 and 1 apply to foreground and background lineages and include sites under purifying selection ($0 < \omega < 1$) and neutral sites ($\omega = 1$), respectively. Site class 2 allows a proportion of positively selected sites ($\omega > 1$) in the foreground lineages, where 2a includes sites under purifying selection ($0 < \omega < 1$) in the background lineages and 2b includes neutral sites ($\omega = 1$) in the background lineages. $2\Delta ln = 30$, d.f. =1, $p < .00001$

[2] Lineages known to eat snakes and/or to be resistant to snake venom (*Erinaceus* sp., and *Mellivora capensis, Sus scrofa) as well as Herpestidae Viverridae, and Eupleridae* were included in the foreground class.

**Supplementary Table 3**. Species that interact with snakes and other venomous species. Asterisk indicates that snakes are venomous

| Mammal Taxa | Nature of Interaction | References |
|---|---|---|
| **Viverridae** | | |
| *Arctogalidia trivirgata* | Eaten by snakes | Eaton et al., 2010 |
| *Civettictis civetta* | Eats scorpions*, aloof to snakes | Schliemann 1990 (?), Amiard et al. 2016, anecdotal; (https://www.youtube.com/watch?v=urIAGGbxGeE&ab_channel=Lala%27slifestyle), https://www.youtube.com/watch?v=frKW9PTYQ7w&ab_channel=ContentMint |
| *Genetta genetta* | Eats snakes, Eaten by snakes | Smithers 1983 (Anecdotal https://www.youtube.com/watch?v=ANx9nvK-pRE&ab_channel=LionMountainTV ) |
| *Genetta tigrina* | Eaten by snakes | Smithers 1983 |
| *Paradoxurus hermaphroditus* | Eats snakes | Grzimek et al., 2004 |
| *Viverra tangalunga* | Eats snakes | Colon et al., 2012, Nowak and Paradiso 1983 |
| **Herpestidae** | | |
| *Bdeogale crassicauda* | Eats snakes* | Sale and Mark 1970 |
| *Bdeogale nigripes* | Eats snakes* | Ray and Sunquist 2001 |
| *Crossarchus obscurus* | Eats snakes* | Struhsaker and McKey 1975, Goldman b 1987, Grzimek et al., 2004 |
| *Cynictis penicillata* | Eats snakes* | Avenant and Nel 1992 |
| *Helogale hirtula* | | Kingdon 1988 |
| *Herpestes edwardsii* | Eats snakes* | Hinton and Dunn 1967 |

| | | |
|---|---|---|
| *Herpestes javanicus* | Eats snakes* | Williams 1918, Tanaka and Mori 2000, Tvrtkovic and Krystufek 1990 |
| *Herpestes ichneumon* | Eats snakes* | Angelici et al., 2005, Ben-Yaacov and Yom-Tov 1983, Stuart 1983, and Delibes et al., 1984 |
| *Galerella pulverulenta* | Eats snakes* | Cavallini and Nel 1990, Fitzsimons 1919 |
| *Galerella sanguinea* | Eats snakes* | Smithers 1983 |
| *Mungos mungo* | Eats scorpions* | Hinton and Dunn 1967, Barett et al. 2012 |
| *Suricata suricatta* | Cooperative mobbing, Eats Scorpions* | Gutzmann et al., 2009., van Staaden 1994, Barett et al. 2012 |
| *Ichneumia albicauda* | Eats snakes* | Smithers 1983 |
| *Paracynictis selousi* | Eats snakes* | Smithers 1983 |

**Supplementary Table 3.** Table of accession numbers for species used in comparative analyses.

| **Accession** | **Species** |
|---|---|
| XM_015086448.2 | *Acinonyx jubatus* |
| XM_002920237.4 | *Ailuropoda melanoleuca* |
| XM_021671790.1 | *Aotus nancymaae* |
| XM_037155882.1 | *Artibeus jamaicensis* |
| XM_034495063.1 | *Arvicanthis niloticus* |
| XM_038337496.1 | *Arvicola amphibius* |
| XM_007183300.1 | *Balaenoptera acutorostrata scammoni* |
| XM_036856972.1 | *Balaenoptera musculus* |
| XM_010859806.1 | *Bison bison bison* |
| XM_019970559.1 | *Bos indicus* |
| XM_005902541.2 | *Bos mutus* |
| NM_176664.2 | *Bos taurus* |
| XM_006073584.2 | *Bubalus bubalis* |
| XM_002749322.4 | *Callithrix jacchus* |
| XM_025887738.1 | *Callorhinus ursinus* |
| XM_010960714.1 | *Camelus bactrianus* |
| XM_031451688.1 | *Camelus dromedarius* |
| XM_032479722.1 | *Camelus ferus* |
| XM_025460248.2 | *Canis lupus dingo* |
| NM_001003144.2 | *Canis lupus familiaris* |
| XM_005675941.3 | *Capra hircus* |
| XM_008062643.2 | *Carlito syrichta* |
| XM_020168889.1 | *Castor canadensis* |
| XM_003478585.3 | *Cavia porcellus* |

| | |
|---|---|
| XM_017538259.2 | *Cebus imitator* |
| XM_014784637.1 | *Ceratotherium simum simum* |
| XM_012047048.1 | *Cercocebus atys* |
| XM_043488325.1 | *Cervus canadensis* |
| XM_043894891.1 | *Cervus elaphus* |
| XM_005393183.2 | *Chinchilla lanigera* |
| XM_007965408.2 | *Chlorocebus sabaeus* |
| XM_037849572.1 | *Choloepus didactylus* |
| XM_006866658.1 | *Chrysochloris asiatica* |
| XM_011943987.1 | *Colobus angolensis palliatus* |
| XM_004674514.2 | *Condylura cristata* |
| XM_003498328.5 | *Cricetulus griseus* |
| XM_004476894.2 | *Dasypus novemcinctus* |
| XM_022570192.1 | *Delphinapterus leucas* |
| XM_024561551.1 | *Desmodus rotundus* |
| XM_013028276.1 | *Dipodomys ordii* |
| XM_004701435.1 | *Echinops telfairi* |
| XM_006878818.1 | *Elephantulus edwardii* |
| XM_022494665.1 | *Enhydra lutris kenyoni* |
| XM_008138537.2 | *Eptesicus fuscus* |
| XM_014851650.1 | *Equus asinus* |
| XM_001499557.6 | *Equus caballus* |
| XM_008528277.1 | *Equus przewalskii* |
| XM_046659125.1 | *Equus quagga* |
| XM_007537522.1 | *Erinaceus europaeus* |
| XM_028124636.1 | *Eumetopias jubatus* |
| XM_003990883.5 | *Felis catus* |
| XM_010607036.2 | *Fukomys damarensis* |
| XM_008568390.1 | *Galeopterus variegatus* |
| XM_030851531.1 | *Globicephala melas* |
| XM_004032824.3 | *Gorilla gorilla gorilla* |
| XM_028762996.1 | *Grammomys surdaster* |
| XM_036075787.1 | *Halichoerus grypus* |
| M93639.1:160-2 | *Herpestes ichneumon* |
| XM_004857704.2 | *Heterocephalus glaber* |
| XM_019653697.1 | *Hipposideros armiger* |
| NM_000079.4 Ho | *Homo sapiens* |
| XM_039239220.1 | *Hyaena hyaena* |
| XM_032753380.1 | *Hylobates moloch* |
| XM_040294499.1 | *Ictidomys tridecemlineatus* |
| XM_004660327.2 | *Jaculus jaculus* |
| XM_027127844.1 | *Lagenorhynchus obliquidens* |
| XM_031021070.1 | *Leptonychotes weddellii* |

| | |
|---|---|
| XM_007451071.1 | *Lipotes vexillifer* |
| XM_032855930.1 | *Lontra canadensis* |
| XM_023542827.1 | *Loxodonta africana* |
| XM_047722226.1 | *Lutra lutra* |
| XM_032594115.1 | *Lynx canadensis* |
| XM_015432377.1 | *Macaca fascicularis* |
| XM_001091711.4 | *Macaca mulatta* |
| XM_011743926.1 | *Macaca nemestrina* |
| XM_011991878.1 | *Mandrillus leucophaeus* |
| XM_017657562.2 | *Manis javanica* |
| XM_036876963.1 | *Manis pentadactyla* |
| XM_027946148.1 | *Marmota flaviventris* |
| XM_015479340.1 | *Marmota marmota marmota* |
| XM_031373391.1 | *Mastomys coucha* |
| XM_046018549.1 | *Meles meles* |
| KR477827.1 | *Mellivora capensis* |
| XM_040745600.1 | *Mesocricetus auratus* |
| XM_012753433.2 | *Microcebus murinus* |
| XM_005346582.2 | *Microtus ochrogaster* |
| XM_041639709.1 | *Microtus oregoni* |
| XM_016206380.1 | *Miniopterus natalensis* |
| XM_035007721.1 | *Mirounga leonina* |
| XM_036254245.1 | *Molossus molossus* |
| XM_001376625.4 | *Monodelphis domestica* |
| XM_029225643.1 | *Monodon monoceros* |
| XM_021152646.1 | *Mus caroli* |
| NM_007389.5 | *Mus musculus* |
| XM_021193505.1 | *Mus pahari* |
| XM_032355993.1 | *Mustela erminea* |
| XM_004774378.2 | *Mustela putorius furo* |
| XM_005857602.2 | *Myotis brandtii* |
| XM_006760599.2 | *Myotis davidii* |
| XM_006083102.3 | *Myotis lucifugus* |
| XM_036320699.1 | *Myotis myotis* |
| XM_008855297.2 | *Nannospalax galili* |
| XM_021703076.1 | *Neomonachus schauinslandi* |
| XM_024732840.1 | *Neophocaena asiaeorientalis asiaeorientalis* |
| KR477835.1:644 | *Neovison vison* |
| XM_030803076.1 | *Nomascus leucogenys* |
| XM_040972098.1 | *Ochotona curzoniae* |
| XM_004577044.1 | *Ochotona princeps* |
| XM_004634743.2 | *Octodon degus* |
| XM_004408997.1 | *Odobenus rosmarus divergens* |

| | |
|---|---|
| XM_020889100.1 | *Odocoileus virginianus texanus* |
| XM_036184821.1 | *Onychomys torridus* |
| XM_033423481.1 | *Orcinus orca* |
| XM_001514832.5 | *Ornithorhynchus anatinus* |
| XM_007940110.2 | *Orycteropus afer afer* |
| XM_008258768.2 | *Oryctolagus cuniculus* |
| XM_040228197.1 | *Oryx dammah* |
| XM_003791505.1 | *Otolemur garnettii* |
| XM_004004575.5 | *Ovis aries* |
| XM_034954553.1 | *Pan paniscus* |
| XM_016950066.1 | *Pan troglodytes* |
| XM_019460101.1 | *Panthera pardus* |
| XM_015541212.1 | *Panthera tigris altaica* |
| XM_021923727.2 | *Papio anubis* |
| XM_037204883.1 | *Peromyscus leucopus* |
| XM_006972439.1 | *Peromyscus maniculatus bairdii* |
| XM_047770453.1 | *Phacochoerus africanus* |
| XM_020990832.1 | *Phascolarctos cinereus* |
| XM_032426208.1 | *Phoca vitulina* |
| XM_032637674.1 | *Phocoena sinus* |
| XM_028509758.2 | *Phyllostomus discolor* |
| XM_007113767.2 | *Physeter catodon* |
| XM_023185999.3 | *Piliocolobus tephrosceles* |
| XM_036422141.2 | *Pipistrellus kuhlii* |
| XM_003776025.2 | *Pongo abelii* |
| XM_043577090.1 | *Prionailurus bengalensis* |
| XM_047871785.1 | *Prionailurus viverrinus* |
| XM_012650725.1 | *Propithecus coquereli* |
| XM_006921218.1 | *Pteropus alecto* |
| XM_039855257.1 | *Pteropus giganteus* |
| XM_011357247.1 | *Pteropus vampyrus* |
| XM_025934934.1 | *Puma concolor* |
| XM_040475435.1 | *Puma yagouaroundi* |
| NM_024485.2 | *Rattus norvegicus* |
| XM_032901959.1 | *Rattus rattus* |
| XM_033111421.1 | *Rhinolophus ferrumequinum* |
| XM_019727497.1 | *Rhinolophus sinicus* |
| XM_017856956.1 | *Rhinopithecus bieti* |
| XM_010353720.2 | *Rhinopithecus roxellana* |
| XM_016134779.2 | *Rousettus aegyptiacus* |
| XM_003921848.2 | *Saimiri boliviensis boliviensis* |
| XM_032292901.1 | *Sapajus apella* |
| XM_003763981.3 | *Sarcophilus harrisii* |

| | |
|---|---|
| XM_004601170.1 | *Sorex araneus* |
| XM_037044418.1 | *Sturnira hondurensis* |
| XM_029932975.1 | *Suricata suricatta* |
| AF540390.1 | *Sus scrofa* |
| XM_038751092.1 | *Tachyglossus aculeatus* |
| XM_037522117.1 | *Talpa occidentalis* |
| KR477838.1 | *Taxidea taxus* |
| XM_025404943.1 | *Theropithecus gelada* |
| XM_033231070.1 | *Trachypithecus francoisi* |
| XM_004375490.3 | *Trichechus manatus latirostris* |
| XM_036746594.1 | *Trichosurus vulpecula* |
| XM_006151116.1 | *Tupaia chinensis* |
| XM_019922603.2 | *Tursiops truncatus* |
| XM_026405652.1 | *Urocitellus parryii* |
| XM_026492523.1 | *Ursus arctos horribilis* |
| XM_008687156.2 | *Ursus maritimus* |
| XM_006204878.2 | *Vicugna pacos* |
| XM_027866151.1 | *Vombatus ursinus* |
| XM_041722809.1 | *Vulpes lagopus* |
| XM_025994006.1 | *Vulpes vulpes* |
| XM_027589625.2 | *Zalophus californianus* |
